# Abundant and metabolically flexible lineages within the SAR324 and gammaproteobacteria dominate the potential for rubisco-mediated carbon fixation in the dark ocean

**DOI:** 10.1101/2024.05.09.593449

**Authors:** Alexander L. Jaffe, Rebecca S. R. Salcedo, Anne E. Dekas

## Abstract

**Background:** Rubisco is among the most abundant enzymes on Earth and is a critical conduit for inorganic carbon into the biosphere. Despite this, the full extent of rubisco diversity and the biology of organisms that employ it for carbon fixation are still emerging, particularly in unlit ecosystems like the deep sea.

**Results:** Here, we generate fifteen deeply-sequenced metagenomes along a spatially-resolved transect off the California coast, and combine them with globally-distributed public data to examine the diversity, distribution, and metabolic features of rubisco-encoding organisms from the dark water column. Organisms with rubisco were detected in the vast majority of all samples, spanning over 1,000 species groups and together comprising up to ∼20% of the total microbial community. At 150 meters and below, potential for carbon fixation via rubisco was dominated by just two orders of gammaproteobacteria and SAR324, encoding either the form I or II rubisco. Many of these organisms also possessed genes for the oxidation of reduced sulfur compounds, which may energetically support carbon fixation. Transcriptomic profiling in the epi- and mesopelagic suggested that all major forms of rubisco can be highly expressed in the deep water column, but are not done so constitutively, consistent with metabolic flexibility.

**Conclusion:** Our results demonstrate that the genetic potential to fix carbon via rubisco is significant and spatially widespread in the dark ocean. We identify several rubisco-encoding species groups that are particularly abundant and cosmopolitan, highlighting the key role they may play in deep-sea chemoautotrophy and the global marine carbon cycle.

## Background

Ribulose 1,5-bisphosphate carboxylase/oxygenase (rubisco) is the key enzyme underlying carbon fixation via the Calvin Benson-Bassham (CBB) cycle and is responsible for the majority of carbon fixation on Earth [1,2]. Employed by plants and photosynthetic phytoplankton, light-driven rubisco is estimated to fix as much as 258 billion tons of carbon dioxide annually [3], which then enters the biosphere as organic carbon. Due to its undeniable importance in photoautotrophy, studies of rubisco have historically focused on organisms from sunlit biomes. However, rubisco is not itself dependent on light, and carbon fixation via rubisco can instead be coupled to chemical energy sources in complete darkness (chemoautotrophy). To date, though, much less is known about the organisms mediating this coupling and their impact on local and global carbon cycling compared to their photoautotrophic counterparts.

One particularly significant dark ecosystem is the deep ocean water column (≥200m water depth), where sunlight is low or entirely absent. There, diverse, abundant, and active microbial communities [4,5] shape global biogeochemistry by remineralizing organic matter exported from the sunlit zone and coupling chemical energy to inorganic carbon fixation to form biomass [6]. In some regions of the deep sea, the biomass resulting from new fixation may reach a similar magnitude to that derived from heterotrophic degradation of organic compounds [7,8], thus representing an important mechanism by which cells’ overall carbon demand might be met [9]. In this way, dark carbon fixation may also represent a significant carbon dioxide sink with potential ramifications for global climate.

For some time, nitrifying archaea fixing carbon with the 3-hydroxypropionate/4-hydroxybutyrate (3HP-4HB) cycle were considered primary drivers of deep-sea chemoautotrophy [10–12]. More recently, single-cell analysis and metagenomic surveys have indicated that lineages using other carbon fixation pathways - including the CBB [5,13,14] - might also be widespread, and contribute more to dark carbon fixation than nitrifying archaea using the 3HP-4HB cycle [15]. However, despite its potential importance in shaping deep-sea biogeochemistry, the full extent of rubisco diversity in the deep sea and the ecological role of organisms that employ it for carbon fixation - including their abundance, distribution, and metabolic strategies - are poorly understood [16]. Additionally, how different forms of rubisco (e.g. form I, II, and III) are differentially distributed and expressed over the physical and chemical gradients of the dark water column has not been carefully explored. Of particular interest is the extent to which form I and form II rubiscos, both of which function in the CBB cycle but are considered to possess different oxygen niches [17], spatially overlap along geochemical gradients in the deep sea.

Here, we examine the diversity, distribution, and metabolism of rubisco-encoding organisms (hereafter termed REOs) in the global ocean, from the epipelagic to the abyssopelagic. Critically, we take a genome-resolved lens to REO diversity, enabling clearer resolution of the taxonomic affiliation, co-occurring genetic characteristics, and abundance compared to gene-centric approaches. Through this lens, we discover over one thousand species groups of bacteria and archaea with variable forms of rubisco and use their genomes to probe changes in carbon fixation potential with depth, both across a spatially-resolved transect in the Northeastern Pacific as well as the global ocean. Supporting our metagenomic analyses, we also leverage public metatranscriptomes that confirm rubisco expression by diverse organisms from the surface through the mesopelagic. Together, our analyses illuminate new aspects of the REO biology in the deep sea and provide foundational information that could help refine biogeochemical models of this vast habitat.

## Results

### Recovery of novel REOs from a deeply-sequenced water column transect

We began our analysis by collecting and deeply sequencing 15 metagenomes (average ∼52 gigabases/sample) from a spatially-resolved water column transect from the California Coast (OC1703A) complementing 13 previously-reported metagenomes from the same expedition (Arandia-Gorostidi et al., *in rev.).* The combined 28-metagenome dataset now spans six sites across nearly 300 km, covering 50 to 4000 m water depth. Paired physicochemical data on these sites is available in Arandia-Gorostidi et al. *in rev.* and Arandia-Gorostidi et al. 2023 [4]. Importantly, this dataset adds significantly to existing sequencing efforts from the dark ocean, especially within the bathypelagic (1000 to 4000 meters depth) where little sampling has been performed to date [18]. Additionally, the depth of sequencing, spatial density of sampling, and paired metadata distinguish it from previous efforts. Here, we utilized the combined dataset of 28 metagenomes to resolve metagenome-assembled genomes (MAGs) - hereafter referred to as the OC1703A MAG set - that accounted for 11.2 to 55.4% (mean ∼41%) of raw metagenomic reads (Table S1). We searched these MAGs for rubisco genes, identifying and subsequently curating medium-to-high quality genome information for 29 REO species groups, ∼76% of which were uniquely assembled at our site and not present in existing public databases (Table S2).

### Taxonomic diversity of rubisco-encoding organisms in the global ocean

We next combined REO MAGs from the OC1703A set with other genomes extracted from existing marine databases [18–22], which were similarly searched at scale for the presence of enzymes in the rubisco superfamily (Methods). The combined set of global REO genomes - encompassing MAGs, single–amplified genomes, and isolates - was subjected to quality filtering (≥50% completeness, ≤10% redundancy) and de-replication, forming ∼1100 clusters at 95% average nucleotide identity, hereafter referred to as ‘species groups’ (Table S2). Next, we sorted the rubisco sequences present in REO species groups into previously described ‘forms’ [23–25] and stringently curated the taxonomy of rubisco-encoding contigs to avoid incorrect inferences caused by metagenomic mis-binning. Curated results included a vast array of rubiscos spanning nearly all known phylogenetic forms, including the form I, form II (Fig. 1ab), and numerous form III-related enzymes (Table S3).

Organisms with the form I were particularly taxonomically diverse, encompassing over 40 orders in 10 non-Cyanobacterial lineages (phyla/classes) (Fig. 1a). The most diverse group of marine REOs were the Gammaproteobacteria, accounting for 125 distinct species groups with the form I enzyme (Table S2). These lineages include some already well-known to perform carbon fixation via rubisco - including the Arenicellales [14] and the PS1 (including members of the SUP05 clade) [26] - as well as numerous newly-reported orders, including two species groups with no current order-level designation in GTDB (order: novel) (Table S2, Fig. 1a). Though without genome-level taxonomy, preliminary searches of a conserved ribosomal marker protein suggested these genomes fall within the described gammaproteobacterial family *Competibacteraceae.* We also recovered evidence for form I enzymes in SAR324 (11 species groups), various Alphaproteobacteria, and the Chloroflexi (Table S2).

**Figure 1.**
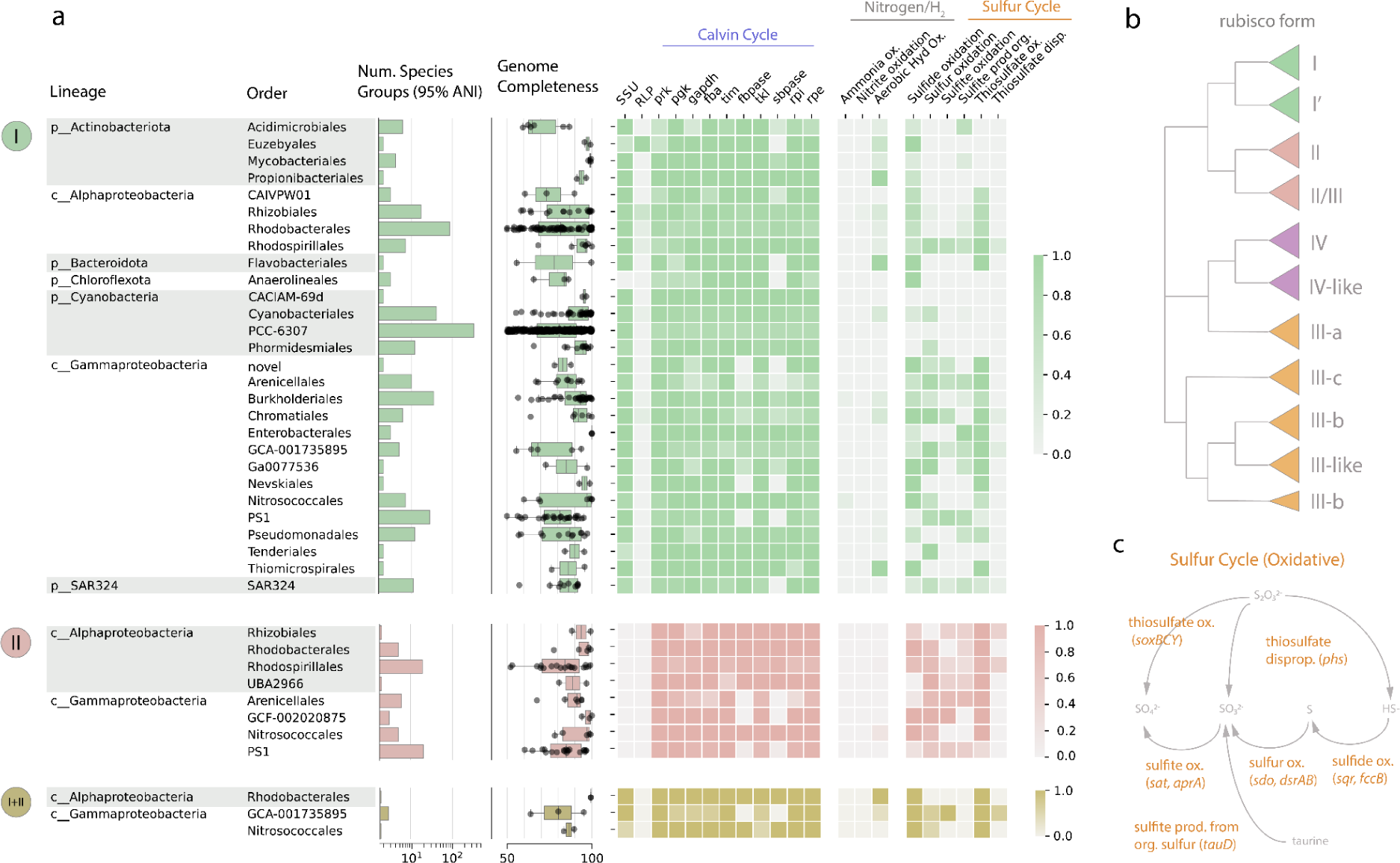
Characteristics of global rubisco-encoding organisms (REOs) with the form I, form II, or both enzyme types (information on form III enzymes can be found in Fig. S1). Organisms were clustered into species groups (95% ANI) and were aggregated at the order level. **a)** For each order-level lineage, the number of distinct species groups, the completeness of representative genomes for each species group, and fraction of representative genomes that encoded the rubisco small subunit (SSU), at least one rubisco-like protein (RLP), CBB cycle genes, and genes involved in oxidation of sulfur or nitrogen compounds for energy gain. Only those order-level lineages with more than one species group are displayed. **b)** Schematic of phylogenetic relationships within the rubisco superfamily adapted from [24]. **c)** Pathway diagram for oxidative reactions involving sulfur, adapted from [93]. Abbreviations: prk, phosphoribulokinase; pgk, phosphoglycerate kinase; gapdh, glyceraldehyde 3-phosphate dehydrogenase; fba, fructose-1,6-bisphosphate aldolase, tim, triosephosphate isomerase; fbpase, fructose-1,6-bisphosphatase; tkl, transketolase; sbpase, sedoheptulose- 1,7-bisphosphatase; rpi, ribose-5-phosphate isomerase; rpe, ribulose 5-phosphate 3-epimerase; ox., oxidation; prod., production, disp., disproportionation; org., organic; Hyd, hydrogen.

In contrast, the diversity of organisms bearing the form II gene, also associated with the CBB cycle, was restricted to a smaller number of groups, primarily in the Alpha- and Gammaproteobacteria (Fig. 1a). Intriguingly, almost all lineages with the form II also contained species groups with the form I, suggesting significant intra-lineage variability in rubisco type in these cases. Furthermore, several proteobacterial lineages - including the Rhodobacterales, the GCA-001735895, and Nitrosococcales (Table S2) - contained a small number of species groups encoding both the form I and form II in the same genome, as has been observed for some autotrophic proteobacteria [17]. Lastly, organisms encoding various form III-related enzymes were recovered in Archaea - including the Nitrososphaerales (Thaumarchaeota) [27,28] - and the Candidate Phyla bacteria [24,29] (Fig. S1, Table S2). Protein sequences for all ∼4400 marine rubisco reported here (609 clusters at 95% amino acid identity) are reported in Table S3.

### Rubisco-associated genes and metabolic features of REOs

Across the rubisco superfamily, there exists significant diversity both in pathway affiliation and genomic configuration (e.g. [24,30]). To examine the metabolic context for these marine rubiscos and assign likely function, we searched representative genomes picked for each global species group for associated genes (Table S4, Methods). As anticipated, only genomes with the form I rubisco encoded the small subunit rubisco (Fig. 1a), which forms a complex with the large subunit and augments its kinetic properties [31]. One exception to this trend was the Chloroflexi, some of whose members encode a divergent form I rubisco that does not require the small subunit to assemble [32]. Other REO lineages without consistent presence of the small subunit are likely due to genome incompleteness, though we cannot rule out the possibility of alternate pathway configurations in all cases.

We consistently recovered the remaining genes in the CBB cycle (Table S5) – the canonical pathway affiliated with the form I and form II rubisco – among the REO lineages, confirming that these rubiscos likely act in functional carbon fixation pathways (Fig. 1a). Importantly, the phosphoribulokinase gene (*prk*), which regenerates the substrate for rubisco so that additional carbon dioxide can be fixed, was nearly omnipresent. On the contrary, fructose-1,6-bisphosphatase (*fbpase*) and sedoheptulose-1,7-bisphosphatase (*sbpase*) were more patchily distributed among the lineages surveyed, including some where representative genomes attained high completeness values (Fig. 1a). This finding suggests that some marine REOs generally may encode streamlined CBB cycles that rely on bifunctional enzymes to cover inner reaction branches, as has been previously described for other organisms [33], or are otherwise reconfigured. Form II/III and III-related enzymes generally co-occurred with genes (*deo and* ribp_isomerase) known to function in an non-autotrophic pathway incorporating CO_2_ with a scavenged nucleotide [34] and lacked the key CBB gene *prk* (Fig. S1). Exceptions to this trend included several methanogenic lineages and one lineage of CPR bacteria that employ *prk* in validated or proposed variants of the CBB cycle (Fig. S1), though it is currently unknown whether these pathways permit autotrophic growth [24,30].

REOs in the deep sea, where light does not penetrate, must power their CBB cycle via chemical energy instead of light energy. Thus, we investigated potential sources of chemical energy by searching genomes for genes involved in nitrogen, sulfur, and hydrogen oxidation (Table S4). Genes involved in oxidation of nitrite and ammonia were rare, and only co-occurred with form I rubisco in six species groups in the proteobacterial orders Burkholderiales, Nitrosococcales, Nitrococcales, and Rhizobiales (Table S6). In contrast, we observed that genes involved in the oxidation of reduced sulfur compounds frequently co-occurred with form I and form II rubiscos, particularly in the Alpha- and Gammaproteobacteria (Fig. 1ac). The genetic capacity for thiosulfate oxidation (*soxBCY)* was widespread, detectable in 69% of proteobacterial species groups with the form I, ∼74% with the form II, and ∼86% with both the form I and form II (Table S6, Fig. 1a). We suggest that these estimates are likely conservative, given that relatively lax thresholds for genomic completeness (≥50%) were employed here.

The ubiquitous detection of the capacity to oxidize thiosulfate broadens and highlights recent evidence from a few individual deep-sea lineages [14,35] and indicates that this metabolic strategy may be a central one among REOs in the deep realm. Similarly, many *sox-*encoding lineages also possessed the genetic potential for sulfide oxidation via *sqr* or *fcc,* as well as the oxidation of elemental sulfur by the sulfur dioxygenase (*sdo*) and/or the reverse dissimilatory sulfite reductase (*dsrAB)* (Fig. 1ac), indicating the potential to catalyze multiple sequential transformations of sulfur compounds. On the other hand, genes enabling the oxidation of sulfite to sulfate were rarer, as were genes encoding the production of sulfite from the organic sulfur compound taurine (Table S6), which could serve as an alternative source of sulfur in the oxygenated water column [14,36]. Finally, in lineages where machinery for sulfur oxidation was rare, like the Actinobacteria, we recovered genetic evidence for uptake hydrogenases (forms 1d, 1l, 2a) that could support fixation through the aerobic oxidation of molecular hydrogen [37,38].

### Abundance and distribution of REOs across a coastal California transect

Previous work has identified REOs in the dark ocean at specific depths [5,13]; however, few studies have traced their distribution over physical and chemical gradients. We leveraged the unique degree of spatial resolution in our newly-reported OC1703A transect to address this sampling gap, and assess the abundance of REOs from the surface into the bathypelagic and from the coast to the open ocean. Metagenomic read recruitment to representative genomes picked for all global species groups, followed by stringent filtering of resulting alignments, revealed that organisms with forms I, II, or III-A rubisco were detectable at all depths surveyed from coast to open ocean (∼300 km from shore) (Fig. 2a). Remarkably, we estimate that these organisms routinely comprised upwards of 10% of the microbial community - as measured by percentage of total sequencing coverage of recovered bins - below 200 m (mean 14.5%), reaching a maximum of 19.5% at 3000 m near the continental shelf. This abundance exceeds those observed in the photic zone (50 and 150 m), though poor genome recovery in the 50 m samples likely hinders this comparison (Table S5).

**Figure 2.**
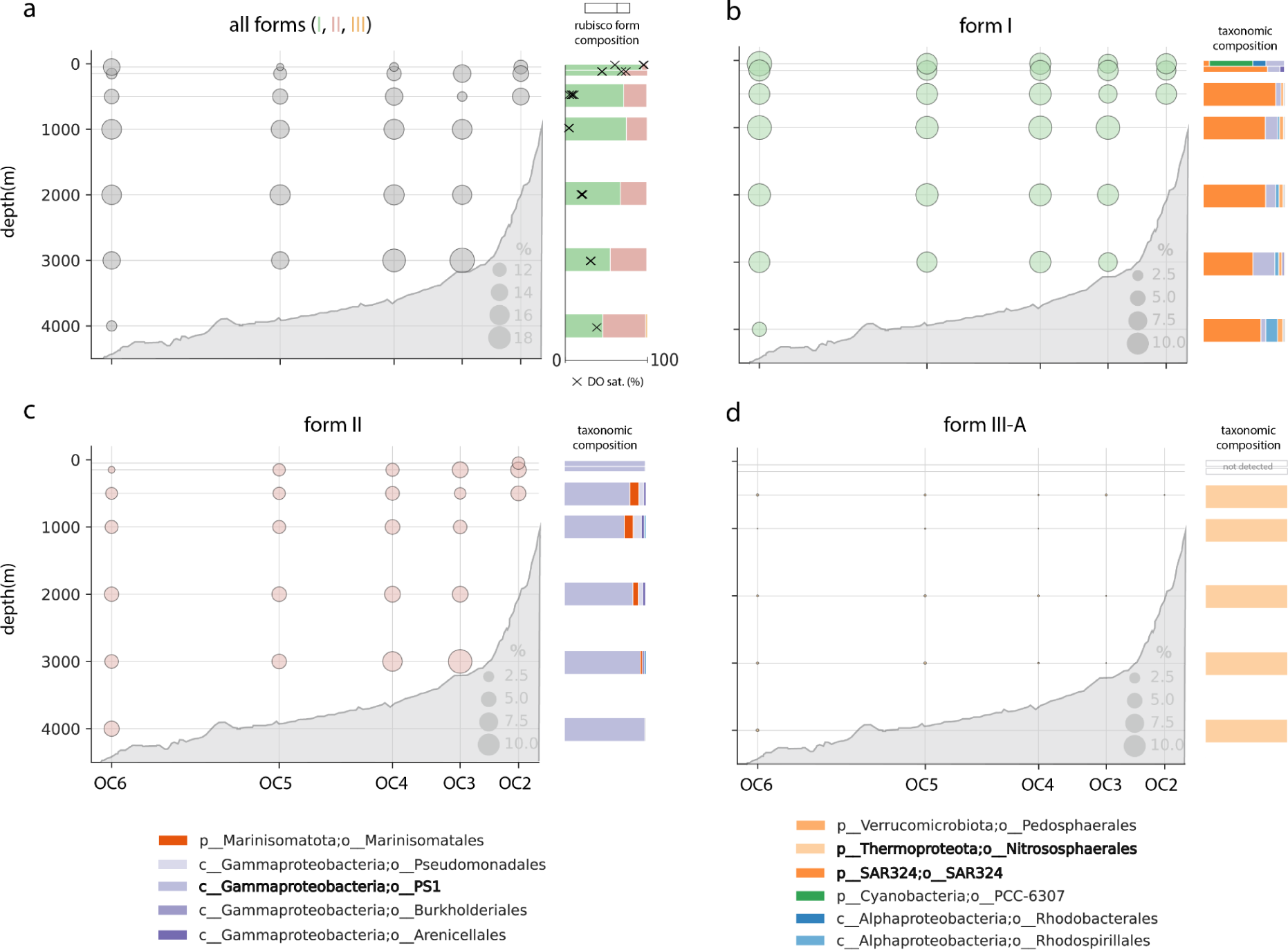
Distribution and abundance of global REOs with different rubisco forms across the OC1703A transect (coastal California). Dotplots represent 5 sites (OC2-6) spanning ∼300 km offshore, sampled across depths. Size of dots represent the relative abundance (calculated as the percentage of total sequencing coverage) of MAGs encoding any form of rubisco, the form I, form II, and form III-A enzyme, respectively. Barplots represent either the composition of rubisco forms (all) or the taxonomic composition of rubisco-encoding organisms with depth (all others). In the upper left panel, percent dissolved oxygen (DO) saturation is overlaid. **N.B.** Seafloor topography was generated by GeoMapApp and is approximate.

While rubiscos at 50 m could primarily be attributed to Cyanobacteria with the form I, we also found evidence for non-phototrophic Alpha- and Gammaproteobacteria at these depths (Fig. 1ab). With the exception of the site nearest to shore (OC2), organisms with the form II rubisco were not recovered at 50 m depth, possibly due to this form’s sensitivity to high oxygen concentrations. On the other hand, at 150 m and below, there was no clear spatial separation of the form I and form II; instead, organisms of both forms co-occurred stably. In fact, we observed that organisms with form II increased proportionally in abundance with depth, despite increasing oxygen concentrations (Fig. 2a). Form III-A rubiscos, attributable to non-ammonia oxidizing Nitrososphaerales - also referred to as heterotrophic marine Thaumarchaeota, or HMT [28] - were present at trace abundances (average ∼0.08% of the community) across the transect at 500 m and below (Fig. 2d).

Intriguingly, REO populations at the OC1703A transect were dominated by a small number of order-level lineages from the global set (Fig. 2). In our transect, organisms affiliated with the SAR324 comprised the majority of the metabolic potential for the form I rubisco (Fig. 2b, side panel). At 100 m and below, organisms from the Gammaproteobacteria PS1 also attained notable abundances; distinct species groups from this same lineage also accounted for the vast majority of form II potential across depth (Fig. 2c). We note that the SAR324 and PS1 lineages were mostly accounted for by singular species groups that were particularly abundant; additional, rarer species were restricted to narrower depth ranges (e.g. epi- or bathypelagic only) (Fig. S2-3). Other order-level lineages, like the Marinistomatales and the Arenicellales, comprised only small portions of form II potential in the mesopelagic, although the latter group can be highly transcriptionally active despite low DNA abundance in some locales [14].

Our genome-resolved approach also allowed us to examine how catabolic pathways supporting carbon fixation varied with depth and space. Along our transect, the metabolic potential for oxidation of various reduced sulfur compounds was widespread throughout the deep water column below 150 m without strong depth patterning - frequently reaching 6-10% of the total binned microbial community (Fig. S4) - mirroring the distribution of metabolically flexible SAR324 and PS1 (Fig. 2bc). At this site, we also observed consistent metabolic potential for sulfite production from organic sulfur compounds, while genes for thiosulfate disproportionation were very rare (<0.1% of the community), but apparently peaked in the bathypelagic around 2000 meters (Fig. S4). On the other hand, REOs with the ability to oxidize ammonia were restricted to the epi- and upper mesopelagic, and we did not detect nitrite or hydrogen oxidizers at the strict thresholds employed here. These findings suggest that reduced nitrogen compounds and molecular hydrogen only minimally support rubisco-mediated carbon fixation at this site, although they likely sustain other, spatially-overlapping types of fixation (e.g. the 3HP/4HB cycle in ammonia-oxidizing Thaumarchaeota ([39], Arandia-Gorostidi et al., *in rev.*).

### Abundance and distribution of REOs across the global ocean

To examine the generality of our results concerning REO abundance and distribution, and explore variations at the global scale, we amassed over 1,000 water column metagenomes from around the world (Fig. 3a, Table S8) and aligned their reads to representative REO genomes compiled above. As before, mappings were stringently filtered on read identity and fraction of genome covered by reads (coverage breadth) to minimize false detections (Methods). As observed in the OC1703A transect, organisms with rubisco were detected in nearly all global samples assessed, whether only deep samples (∼93%), all samples (∼92%), or only autotrophic forms I and II (∼89%) were considered. Where detected, we computed the relative abundance of each REO (Table S9) and summed these values by rubisco form to visualize relationships between abundance and depth (Fig. 3b).

**Figure 3.**
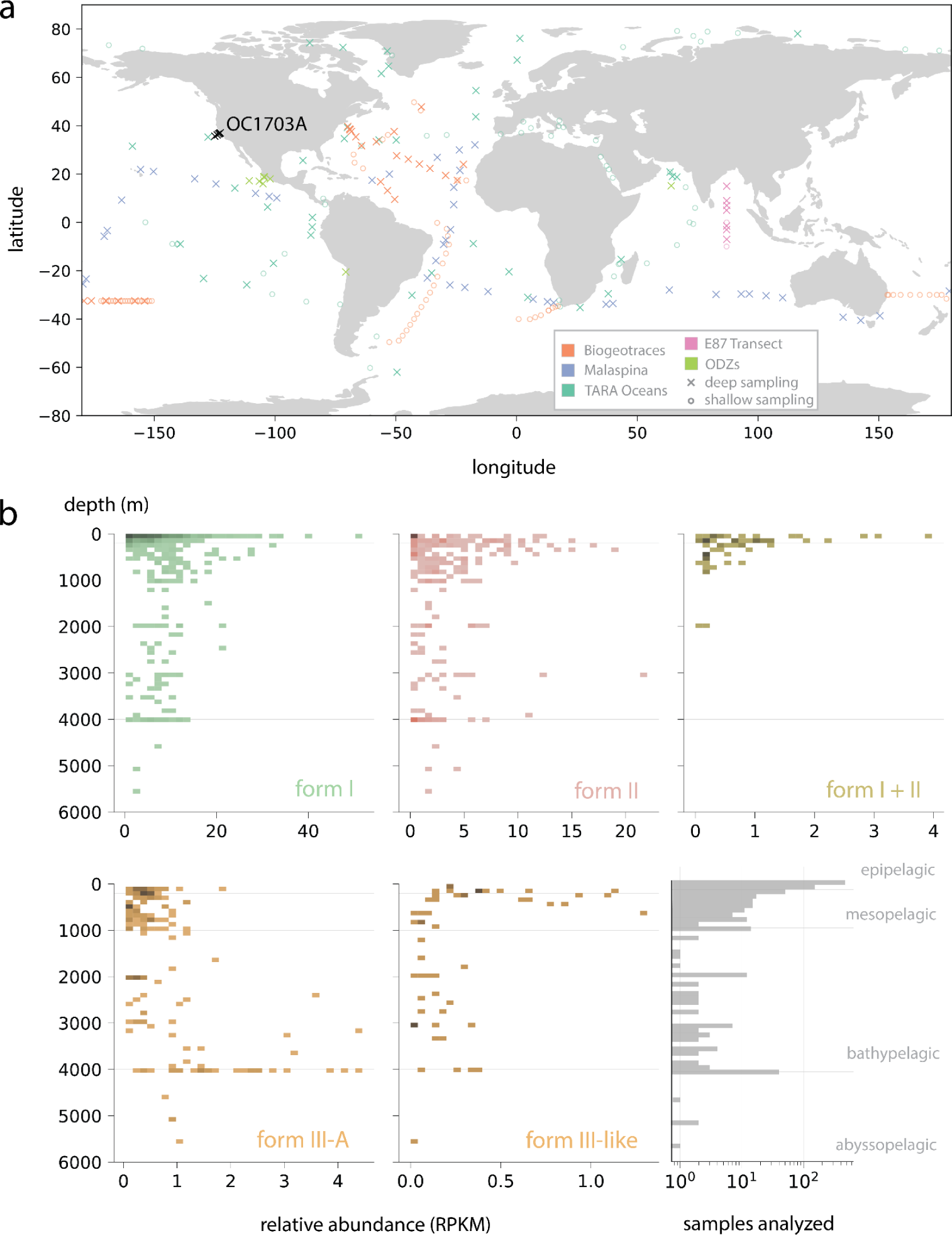
Distribution and abundance of rubisco-encoding organisms (REOs) across the global ocean. **a)** Water column sites where REOs were detected. An ‘X’ refers to a site where at least one deep (≥200m) sample was taken, whereas open circles indicate sites with only shallow (<200m) samples. **b)** Relative abundance of REOs with different rubisco forms expressed as RPKM (reads per kilobase million). Shaded cells indicate the detection of one or more organisms in a given depth/RPKM bin, with hue intensity indicating the density of observations in that bin. Lower rightmost panel describes the number of metagenomes analyzed by depth.

As in the OC1703A transect, we observed that organisms with the form I rubisco attained the highest relative abundances globally of all forms, reaching maximum values (≥40 reads per kilobase million, RPKM) in some epipelagic samples (Fig. 3b, Table S9). The relative abundance of these groups remained fairly constant with depth through to the boundary of the abyssopelagic. Similar patterns were observed for organisms with the form II and III-like (typically CPR bacteria and DPANN archaea), although at a smaller scale (Fig. 3b). On the other hand, lineages encoding both the form I and form II in the same genome were essentially restricted to the mesopelagic, and were rarely observed below 1000 meters. Organisms with the III-A rubisco (generally non-ammonia oxidizing HMT) displayed the reverse pattern, apparently peaking in relative abundance below 3000m (Fig. 3b), consistent with previous reports [28]. Most major rubisco forms were patchily distributed below 4000m, but poor sampling hinders a complete picture in this ocean layer.

Leveraging the global breadth of our metagenomic database, we next investigated the biogeography of individual species groups based on the read mapping described above. To reduce the impacts of uneven spatial sampling, we clustered all metagenomes drawn from 200 meters depth or below into ‘locales’ within 10 kilometer radii (Table S8, Methods) and counted the number of locales in which each REO species group was detected. Intriguingly, most REO species groups were provincial, in other words appearing at only one or a few locales (Fig. 4a). In general, provincial REO species groups attained lower mean abundances (Fig. 4b).

**Figure 4.**
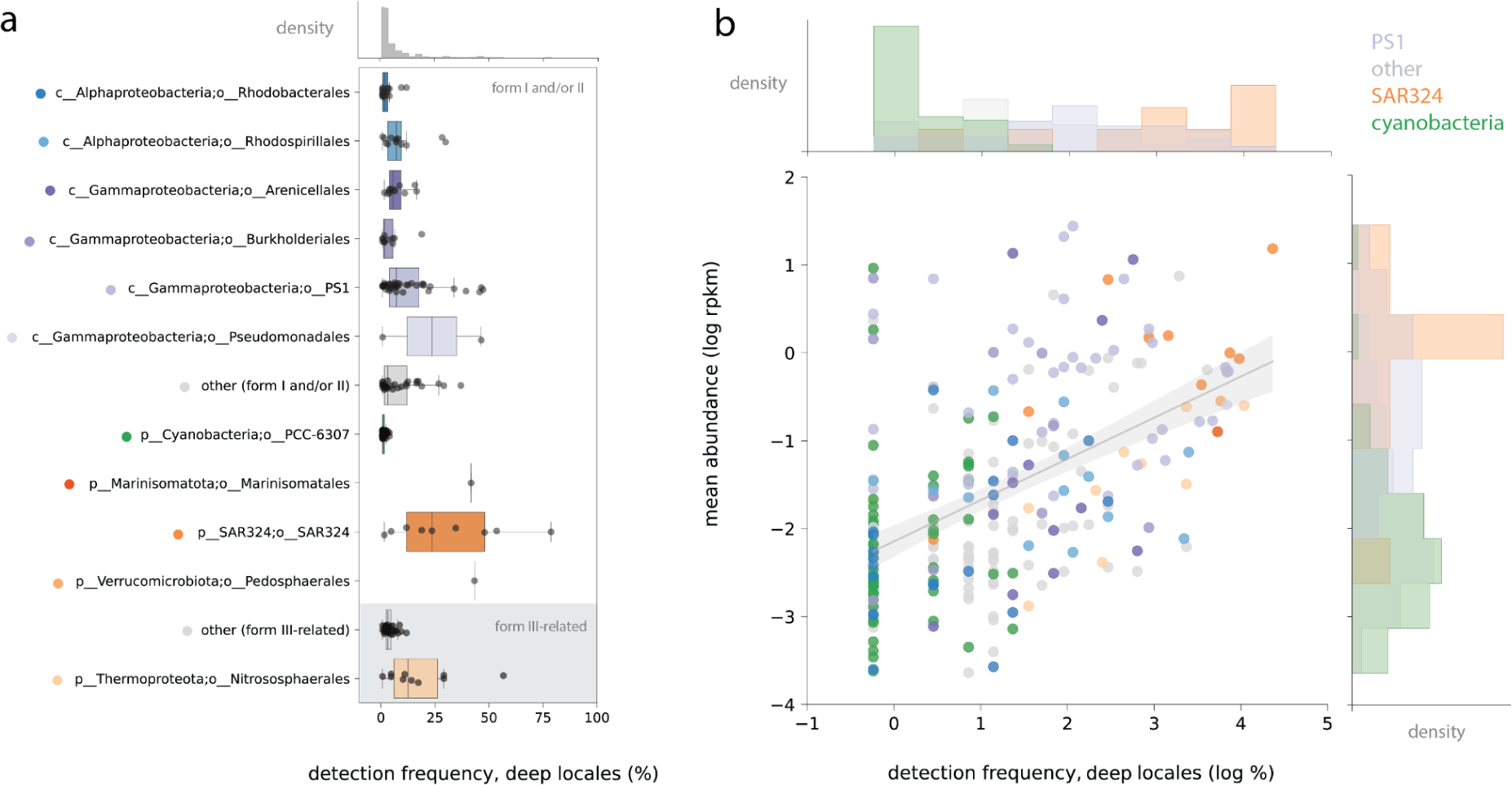
Biogeography of REO species groups from various bacterial and archaeal lineages within the global metagenomic dataset. **a)** Frequency detection of REO species groups among deep water locales (≥200m depth), ordered by form and taxonomy. A subset of lineages are shown, the remaining lineages are marked here as ‘other’. Colors correspond to those in **b),** which depicts frequency of detection (again, only deep locales) plotted against mean relative abundance. The trendline reflects a simple linear model with a 95% confidence interval. In this case, a species group was considered present at a given locale if it was detected in any size fraction, and at any depth ≥200 meters.

We did, however, also recover evidence for multiple more cosmopolitan species groups, where representative genomes were detected in over one third of global deep locales. These species groups largely derived from the same lineages that dominated rubisco form I and II metabolic potential in the OC1703A transect: SAR324, PS1, and to a lesser extent, the Pedosphaerales, Pseudomonadales, and Marinistomatales (Fig. 4a). Remarkably, one SAR324 species group that was dominant at OC1703A (SG114_2) was also detected in almost 80% of global deep-water locales, the highest of any in our dataset. Visualization of SG114_2 abundance with depth over the global dataset showed that this species is generally rare at the ocean surface and increases in abundance towards a peak at or around 2000 meters (Fig. S5), implying that it is a deeper-adapted ecotype. A similar pattern was observed for SG210_1 from the Gammaproteobacterial order PS1 (∼45% of locales), which was the most dominant lineage encoding the form II rubisco in the OC1703A transect (Fig. S3, Fig. S5).

### Patterns of gene expression in REO genomes

While evidence for expression of rubisco has been reported at a number of water column sites, the extent to which this expression is quantitatively important for endemic taxa remains poorly understood. We next examined organism-specific rubisco expression by collecting approximately 200 metatranscriptomes from the world oceans across depth, as gene expression studies of the dark water column are sufficiently rare as to prevent us from focusing on deep datasets exclusively. We aligned sequenced transcripts to the global REO genome set, stringently filtered these alignments such that transcripts could be uniquely ascribed to a single organism, and finally required that genes surpass a minimum threshold of sequencing coverage (coverage breadth ≥50%) to be considered expressed. While we acknowledge that such an approach is likely biased against rarer or less active organisms that obtain lower sequencing coverage, here we focused on those with the clearest signal of gene expression to minimize the impacts of ambiguous mapping and/or trace DNA contamination.

First, among those organisms that we deemed transcriptionally active (at least 50 genes with detectable expression in a sample), we ranked the expression of rubisco relative to all other expressed genes. Broadly, this analysis revealed that diverse REOs actively express the major forms across depth, sometimes within the top 10% of all genes in their genomes (percentile expression ≥90%, Fig. 5a). We observed no major differences between these forms with depth. However, many non-cyanobacterial REO expressing rubisco genes did so at median or lower levels compared to other genes in the genome (percentile expression ≤50%), with overall distribution across depths and forms appearing relatively uniform (Fig. 5a, ‘non-cyanobacteria’ panel). This stood in stark contrast to expression patterns for shallow cyanobacteria, where we observed a radically left-skewed distribution, indicating very high expression of form I rubisco under most conditions. This discrepancy may reflect major differences in rubisco turnover rates or fundamental metabolic strategies between generally autotrophic Cyanobacteria and the bacteria and archaea profiled here. Importantly, we note that in most cases, rubisco expression among active non-cyanobacterial species groups was zero (55% of observations) or non-zero but below threshold for active expression (∼32%) (Table S10). Interestingly, these same REOs did display active rubisco transcription in other samples (including PS1 and SAR324), highlighting dynamic transcriptional profiles for these groups. (Table S10). Thus, our results not only provide further evidence that diverse lineages use rubisco to fix carbon through the deep sea, but also suggest that it may constitute one of multiple anabolic strategies for some hosts.

**Figure 5.**
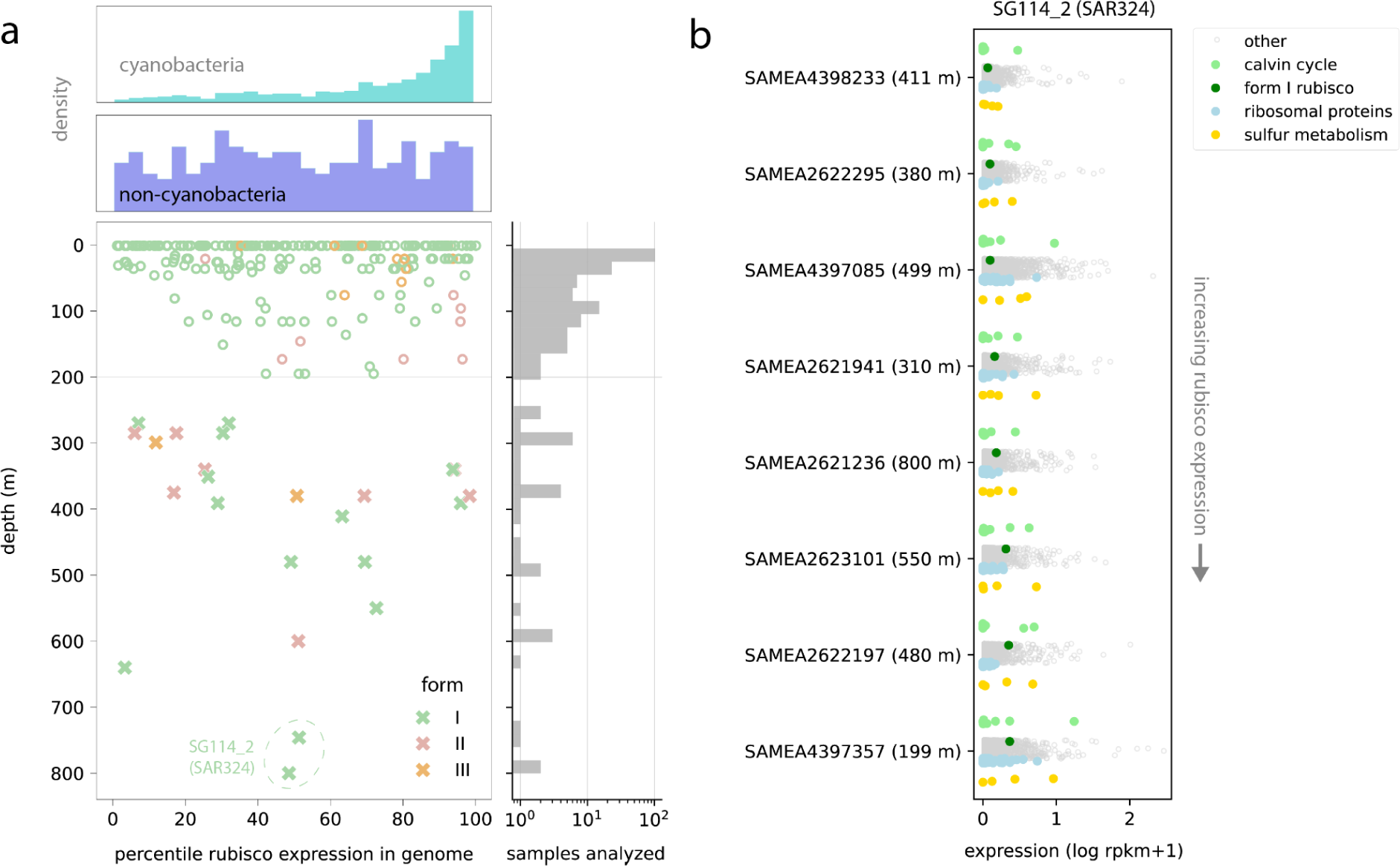
Transcriptional profiles of globally abundant REO. **a)** Extent of rubisco expression in non-cyanobacterial species as a function of depth. Each point represents a transcriptionally active genome in a single sample with above-threshold rubisco gene expression (≥50% gene coverage breadth). X-axis values describe the percentile expression of rubisco among all transcribed genes in that genome (Methods). For clarity, active organisms with rubisco expression that was below the minimum threshold are not displayed. Upper histograms depict the density of observations, with equivalent measurements for cyanobacterial genomes shown for comparison. Rightmost panel describes the number of metatranscriptomes analyzed with depth. **b)** Genome-wide transcriptional profiles for an abundant SAR324 species group (SG114_2) across a subset of water column samples. **N.B.** The position of different gene sets within a sample does not convey quantitative information.

The genome-centric approach employed here also enabled us to scrutinize the expression profiles of individual organisms. Given the prevalence of SAR324 species group 114_2 (SAR324) in our set of deep metagenomes (Fig. 5a), we reasoned that this organism might also be frequently detected in metatranscriptomes, even with the lower extent of global sampling in current databases. Indeed, we found that this species group was transcriptionally active in over half of metatranscriptomes sampled from below 200 meters, and, notably, was responsible for the two deepest observations of rubisco expression in our transcriptome dataset (∼800 meters, Fig. 5a). We observed no relationship between depth and extent of rubisco expression (here, form I), suggesting that other factors likely govern the regulation of carbon fixation machinery in this organism (Fig. 5b). Using expression values computed for all genes in the genome, we next compared rubisco expression to other gene sets of interest. Across samples, we noted concurrent expression of the CBB cycle and sulfur oxidation machinery, consistent with the latter’s potential role in energetically supporting carbon fixation (Fig. 5b). Intriguingly, expression of genes within these pathways displayed high variance, with transcription levels of triosephosphate isomerase (CBB cycle), adenylylsulfate reductase (sulfite oxidation), and *soxB* (thiosulfate oxidation) often surpassing those of the ribosomal proteins (Table S11).

## Discussion

### Diversity, abundance, and distribution

Organisms with rubisco are increasingly recognized to compose significant portions of carbon-fixing communities in the dark ocean [13,14]. Here, using a rubisco-centric screening approach, we significantly expand the number of bacterial and archaeal lineages known to encode this gene and demonstrate that REO species groups are widespread throughout the marine water column. By performing our analyses in a depth-resolved fashion, we show that non-cyanobacterial REO are not only highly abundant – in some cases, reaching nearly a fifth of the total microbial community below 200 m (Fig. 2) – but are also commonly detected in shallow regions where sunlight penetrates (Fig. 2). Co-occurrence with phototrophs in some shallow samples suggests that chemoautotrophy may remain advantageous despite ample availability of organic matter [16] and light energy. Supporting this, we detected transcription of carbon fixation machinery across essentially the entire water column where data has been generated (Fig. 5). Future gene expression studies of the bathypelagic will illuminate the extent to which these trends extend to depths where nutrient limitation intensifies, if technical challenges to producing low biomass RNA libraries can be surpassed. However, transcription of rubisco by at least one REO in sediment traps from 4000 meters [40] is suggestive that microbial carbon fixation via rubisco occurs actively deep into the ocean interior.

Few studies to date have examined the ubiquity of individual chemoautotrophic taxa over the vast expanse of the global ocean. Drawing on the public data, we demonstrate that REO genomes are detected in the vast majority of metagenomic samples from the major basins, spanning functional and phylogenetic forms in the rubisco superfamily (Fig. 3). Among the most ubiquitous REO were members of the SAR324, largely encoding the form I rubisco, as well as members of the PS1 (including the SUP05), which frequently encoded the form II enzyme type. Remarkably, one species from the former lineage was recovered in over three quarters of deep locales analyzed, consistent with a previous study that consistently detected a singular SAR324 species within global ∼4000 meter sites [5]. These combined findings point to cosmopolitan taxa that may anchor community potential for carbon fixation by rubisco, even if less abundant species contribute more to instantaneous transcription under some conditions [14]. Intriguingly, REO genomes with **both** the form I and II enzyme types differed from other major types examined here in that they were barely detected below 1000m (Fig. 3b), despite the metabolic flexibility afforded by this gene inventory [17]. Instead, organisms with multiple rubiscos may be selected for in zones where fluctuations in oxygen/carbon dioxide are greater in magnitude, or might instead be limited by other environmental/biological factors.

Biochemical studies have suggested significantly different kinetic properties for form I and form II enzymes, which like all *bona fide* (form I-III) rubisco can utilize both carbon dioxide and oxygen as substrates. Largely due to decreased ability to discriminate between substrates (*S_c/o_*), form II enzymes are thought to be specialized to lower oxygen, higher CO_2_ niches [17], where competition for enzyme active sites by undesirable oxygen molecules is reduced. Given these properties, we were surprised to observe that organisms encoding either the form I **or** form II enzyme co-occurred through the water column both in the OC1703A transect, and showed no consistent relationship between abundance and oxygen concentration (Fig. 2). To explain this unexpected result, we hypothesize that these REOs might occupy distinct microniches within the dark water column, where they mediate carbon fixation at different rates according to their rubisco gene inventory. Supporting this notion is growing evidence that anaerobic conditions might arise in particle interiors where microbial activity depletes local oxygen levels [41], and that such aggregates may be more widespread in the oxygenated ocean than previously recognized [42]. However, while some REO have been imaged in association with aggregates [13], their genomes do not always contain canonical genes for particle-associated lifestyle (e.g. attachment) [43]. Currently, the predominant lifestyles of deep-sea REOs remains an open question, to be addressed more comprehensively in future research.

### Metabolism: genes and transcriptomes

Energy supporting carbon fixation in marine chemoautotrophs may be derived from a variety of sources, including oxidation of hydrogen, nitrite or ammonia, and reduced sulfur compounds [16,44,45]. Evidence for the particular association of rubisco and genes for oxidation of various reduced sulfur compounds (e.g. sulfide, thiosulfate, sulfite) is accumulating from both bulk gene quantification approaches as well as genome-resolved metagenomics [14,43,46]. Recently, a firmer link between these processes was established by demonstrating that amendment of deep-sea water with thiosulfate stimulated dissolved inorganic carbon fixation up to four times compared to an unamended control [35]. Here, we expand upon this growing body of evidence by reporting a wide distribution of sulfur oxidation genes among the largest set of REO genomes assembled to date. If reduced sulfur compounds can indeed serve as a source of energy supporting carbon fixation, our results suggest that this strategy is likely the primary one employed by REOs in the deep water column. Importantly, at least in the OC1703A transect, we predict that these organisms are spatially widespread across depth (Fig. 1, S4), and thus are unlikely to be solely the result of transport from anoxic coastal regions, as has been proposed for a subset of groups [26]. Alternatively, our results would be consistent with the notion that some REOs localize to suboxic niches where sulfate reduction can occur [13,42,47], or to interfaces where both oxygen and reduced inorganics would be available [17].

Outside of the Cyanobacteria, very few REOs identified in our study are known to function as obligate autotrophs dependent upon chemical/light energy and carbon for growth. In fact, a growing body of literature suggests that individual representatives of major bacterial lineages with rubisco may live mixotrophically, at least transiently depending on organic matter when it is available. Specifically, studies have reported presence of various genes for organic compound transport in genomes also encoding rubisco and sulfur oxidation machinery [5,13,48,49]. Recently, these genomic claims were bolstered by experimental evidence demonstrating tandem uptake of inorganic and organic substrates in the same population of cells [14]. However, to date, the potential for mixotrophy among deep sea REOs has not been systematically examined. While not a facet of our genomic analyses here, we did observe a wide range of rubisco expression values (Fig. 5), and many cases where organisms were active but no rubisco expression was detected. From these results, we infer that some form I and II-bearing species were sampled while growing heterotrophically, as obligate chemoautotrophs likely continually express the CBB cycle. This metabolic flexibility may permit REOs to persist through periods of low organic matter availability in their vast and dilute habitat.

### Conclusions

Overall, our results yield important insights into the diversity, distribution, abundance, and metabolism of widespread lineages with the genetic potential to perform inorganic carbon fixation via rubisco in the ocean interior. The knowledge gained could inform efforts to model the microbial contribution to organic carbon supply/demand in this habitat, which is currently not well resolved [6,9]. One important contribution of this study is the notion that abundant global REOs fall into relatively few taxonomic (e.g. SAR324 and PS1) and metabolic types (e.g. sulfur oxidizers) facilitating their future inclusion in trait-based biogeochemical models. Further, that some individual REOs are found in the majority of marine communities identifies them as attractive targets for cultivation efforts, which could ultimately enable measurement of process rates in addition to the ecological parameters identified in this study.

Of course, in rationalizing the roles of carbon fixers in deep-sea biogeochemistry, it is important to keep in mind other pathways beyond the CBB cycle. This is particularly true in the vast expanses of the ocean that experience transient or permanent anoxia, and thereby may select for alternative pathways with higher oxygen sensitivity and reduced energetic demands [50,51]. While preliminary work from a subset of depths suggests that the CBB cycle is dominant in the dark ocean water column [5], additional study is needed to determine the relative importance of the CBB to other pathways on a larger spatial scale, and relate emergent patterns to a broader slate of physical and chemical parameters.

## Methods

### Metagenomic analysis of microbial communities from the OC1703A transect

Samples for metagenomic analysis were collected in the Pacific Ocean off the USA California coast (north of Monterey Bay) using the *R/V Oceanus* in the spring 2017 and processed as described in Arandia-Gorostidi et al., 2023 [4]. Briefly, Niskin bottles were used to collect seawater at 50, 150, 500, 1000, 2000, 3000, and 4000 meters water depth at 6 sites along a 300 km transect. Seawater (5-25 liters depending on depth) was filtered onto 0.2 µm Sterivex units (Millipore, Germany) and flash frozen in liquid N_2_ before storage at −80°C. DNA extraction was performed using the AllPrep DNA/RNA kit (Qiagen, Valencia, CA, USA). Paired-end metagenomic sequencing was performed on a NovaSeq 6000 S4 platform (150 bp reads) at the UC Davis DNA Sequencing Facility. In this study, we report metagenome information for 3 sites - OC2, OC4, OC5 (15 samples total across depths) – for the first time, in addition to the 2 sites - OC3 and OC6 (13 samples) – reported previously (Arandia-Gorostidi et al., *in rev.*).

To generate a set of MAGs representing the microbial community across sites, sequencing reads were quality-filtered using bbduk (sourceforge.net/projects/bbmap), assembled using MEGAHIT (v1.2.9) [52], and binned using the metabat tool suite [53] as described in Arandia-Gorostidi et al. *in rev.* Cross-mapping was performed for a subset of sample pairs using bowtie2 [54], and mapping results were passed to metabat2 [53] for differential coverage binning. Select output bins were refined using the anvi-refine function from Anvi’o [55], in which member contigs were visualized and manually removed if they displayed abnormal GC content or coverage profiles across the set of metagenomes.

### Construction of a global dataset of REOs from the marine water column

We combined the newly-resolved OC1703A MAGs with genomes drawn from existing collections of metagenome-assembled genomes (MAGs), single-amplified genomes (SAGs), and other genomes from the global ocean - specifically, the Ocean Microbiomics Database [18], the Ocean DNA catalog [19], and several studies targeted low-oxygen ocean regions [21,22,56]. The combined set was subjected to a preliminary screen for rubisco using graftM [57] and a custom package built from rubisco superfamily sequences reported in [23] clustered at 75% identity using usearch [58]. This custom package is available at the project’s GitHub repository (github.com/alexanderjaffe/rubisco-genomics/). To control for false positives among protein hits from graftM, sequences were secondarily annotated using kofamscan [59], and only those hits with top hits to KEGG Orthology numbers K01601, K08965, and K25035 (describing rubiscos and rubisco-like proteins) at e≤1E-5 were retained. Genomes containing putative rubiscos were then assigned a genome-level taxonomy using GTDB-Tk [60], filtered at ≥50% completeness and ≤25% redundancy/contamination as computed by CheckM [61], and clustered at 95% average nucleotide identity using dRep [62].

To account for the possibility of mis-binned rubisco genes, including among genomes drawn from public repositories, we analyzed the taxonomic profile of rubisco-encoding contigs in all genomes (using the method outlined in Arandia-Gorostidi et al. *in rev.*) and compared them to overall genome taxonomy assigned by GTDB-Tk. Briefly, contigs were assigned a taxonomic affiliation based on a consensus of individual protein taxonomies, and manually curated when short or ambiguous. Contigs were flagged as misbinned if consensus taxonomy deviated from genome taxonomy at the phylum level; most frequently, misbinned contigs encoded eukaryotic rubisco sequences that could be easily identified by this methodology. A refined set of REO genomes was created by removing these contigs and any rubisco-encoding scaffold ≤1500 bp in length. Clustering was performed again as above to produce a set of species groups where all members attained ≥50% completeness and ≤10% redundancy.

### Identification and curation of encoded rubisco proteins

Each genome in the quality-filtered set was subjected to gene/protein prediction by Prodigal [63] (*single* mode). Resulting proteins, with the exception of those from contigs encoding rubisco that were either ≤1500 bp in length or misbinned, were combined and searched against a series of custom HMMs describing the major forms within the rubisco superfamily using HMMER [64]. These HMMs were built from multiple sequence alignments constructed from sequences reported in Prywes et al. [23] clustered at 90% identity using usearch [58]. HMM results were parsed using the SearchIO package in Python, and results were filtered to those in which query sequences covered at least 50% of the model. If a given query hit multiple models, the model with the highest HMM score was chosen.

To confirm these preliminary form-level annotations, sequences were also classified via a phylogenetic approach using the graftM package described above. Where HMM-based and phylogenetic classifications differed (∼10% of 609 clusters), classifications were manually curated through a combination of BLAST and phylogenetic tree visualization. Curation was aided by forming sequence clusters at 95% identity using vsearch (*--id 0.95 --maxrejects 0 --maxaccepts 0*) [65]. Finally, each species group was assigned a set of representative rubisco sequences by determining consensus rubisco inventories of its member genomes. This approach allowed us to account for cases in which individual genomes encoded fragmentary rubisco that were lost due to filtering of HMM results. Clusters with no rubisco proteins were removed, yielding a final set of 1070 clusters for downstream analysis.

### Analysis of genome-level metabolism

We further analyzed the metabolism of the genome clusters from above, for each selecting a representative genome of the highest quality using dRep results. Genome clusters encoding only a Form IV rubisco/RLP were excluded from this analysis. For those included, proteins predicted from representative genomes were re-annotated using kofamscan [59]. HMM results were filtered using the provided score thresholds for each model; in some cases, thresholds were relaxed using a previously described method that aims to include divergent or fragmented sequences by visualizing HMM scores in a phylogenetic context [66]. Results for each representative genome were then queried for a set of metabolic genes known to function alongside rubisco in various CO_2_ fixation/incorporation or in pathways that oxidize sulfur and nitrogen compounds for energy gain [67]. Gene names, their associated KEGG accession numbers, and operational thresholds employed here are reported in Table S4. A metabolic pathway was considered present if at least one marker gene was found; however, in the case of sulfite oxidation, we required the presence of at least two (*aprA* and *sat*).

Results were further refined in several cases where bioinformatic identification using HMMs alone was not adequate to discriminate between the gene of interest and closely related homologs. Protein sequences for putative *dsr* (dissimilatory sulfite reductase) genes were merged and compared with a set of curated references [68] using blastp. Putative *dsr* genes with a combined 50% or more percent identity and 70% coverage to known members of the oxidative clade were retained, while any genes most closely matching reductive members were discarded. Similarly, putative sulfur dioxygenases (*sdo)* protein sequences were aligned with a set of references and examined for the presence of two residues found to distinguish true *sdo* from related metallo-β-lactamases [69]. Those sequences without either key residue were discarded. Nitrite oxidoreductases were distinguished from other members of the Type II DMSO reductase family by placing them in a phylogenetic tree with a diverse set of references [70]. Putative ammonia monooxygenases were distinguished from other copper membrane monooxygenases [71] in a similar fashion. In both cases, alignment was performed by MAFFT [72], alignment trimming with trimal (*-gt 0.1)* [73], tree inference with FastTreeMP [74], and tree visualization with iTol [75].

Lastly, the genetic capacity for H_2_ oxidation via was examined by comparing all predicted protein sequences from the representative genome set to published NiFe hydrogenases, using the reference set and annotation technique reported by [38]. Sequences were filtered to those likely permitting energy gain via aerobic oxidation of H_2_ (forms 1d, 1l, and 2a), and these form assignments were confirmed with the above alignment/tree-building approach.

### Abundance and distribution of REOs across the OC1703A transect

The presence of REOs was first quantified across a 28-sample transect from the California coast. As described above, these samples were sequenced, quality-controlled, assembled, and binned, yielding a number of medium-high quality MAGs representing the full microbial community at these sites. This pool of MAGs was added to the global REO set and all genomes were re-clustered at 95% ANI using dRep. Representative genomes for each cluster were chosen using default dRep parameters, unless the cluster included a previously identified REO genome, in which case that genome was selected. If a cluster contained multiple REO genomes, the one with the highest completeness was selected as a representative. To avoid oversampling REOs compared to organisms without rubisco, and thus skewing abundance metrics, only REOs clustering with MAGs assembled from the transect itself were retained.

Next, trimmed metagenomic reads from the 28 samples were mapped to representative genomes using bowtie2 [54] (default parameters). Coverage for each bin was computed using CoverM (https://github.com/wwood/CoverM) with a 95% read identity threshold. Genomes were considered present in a given sample if 50% or more of its bases were covered by reads (coverage breadth). Relative coverage of each genome was calculated by dividing its mean coverage by the summed coverages of all genomes detected in that sample. Abundance patterns for genomes encoding any form of rubisco were then visualized as a function of genome taxonomy or gene inventory using Python.

### Abundance and distribution of REOs across the global ocean

To examine abundance and distribution patterns of REO at a larger scale, we amassed a list of nearly 1,000 water column metagenomes from previous studies [5,20,22,56,76–85] as well as corresponding metadata. Raw reads were downloaded from the Sequence Read Archive, and, if multiple sequencing runs were listed for a single sample, their reads were merged. Next, forward and reverse read files were trimmed using bbduk.sh from BBTools (sourceforge.net/projects/bbmap/) (*ref=adapters ktrim=r k=23 mink=11 hdist=1 tbo qtrim=r trimq=25 minlen=20*). Trimmed reads were then mapped against the non-redundant REO genome set using bowtie2 (default parameters), and the resulting alignment files were stringently filtered using inStrain (*--min_read_ani 0.95 --min_mapq 10*) [86]. A Snakemake [87] workflow implementing these sequential processing steps is available via GitHub (github.com/alexanderjaffe/rubisco-genomics/) and is conceptually visualized in Fig. S6.

Once processed, inStrain results were read into a Pandas dataframe in Python and further filtered on coverage characteristics to reduce the incidence of false detections. Specifically, a genome was considered present in a sample if it attained ≥50% coverage breadth and ≥50% of expected coverage breadth as defined by inStrain. Next, RPKM values were calculated for each genome-sample pair using the number of filtered reads mapping specifically to the genome, the genome length, and the total number of trimmed metagenomic reads in the mapped sample. Genomic RPKM values were summed per sample for all organisms with the same rubisco genes, and subsequently displayed as a function of depth (Figure 3b). In this case, only size fractions corresponding to prokaryotic cells were used (size fractions targeting viral communities were omitted). For ease of visualization, values were sorted into depth bins of 100 meters.

Geographic coordinates for all metagenomic samples analyzed were plotted with GeoPandas in Python. To control for unequal spatial sampling, site coordinates were clustered into ‘locales’ using density-based spatial clustering of applications with noise (DBSCAN) according to a published protocol [88], with an epsilon value of 10 kilometers. To determine frequency of detection of individual REO species across sites, we determined the number of unique locales in which that species was detected at the breadth thresholds described above. A species was considered present if detected in any size fraction or at any depth ≥200 m at a given site.

### Genome-resolved transcriptomics of REOs

Gene expression was assessed in ∼200 publicly available metatranscriptomes from TARA Oceans, as well as several studies of anoxic or oxygen-deficient ocean regions [89–92] (Table S10). Raw reads were first downloaded, trimmed, and mapped against the non-redundant genome set as described above. For each metatranscriptome successfully mapped, read coverage of any genomes detected was quantified using pysam (https://github.com/pysam-developers/pysam). Specifically, we counted the number of reads mapping stringently (*min_mapq 20*, *max_mismatch 5*) to each gene and used this to compute gene-wise mean coverage and breadth of coverage. To reduce computational complexity, only those genomes recruiting 500 reads or more from the transcriptome were subjected to gene-by-gene analysis. A Snakemake workflow detailing this is available via GitHub (github.com/alexanderjaffe/rubisco-genomics/) and drawn out in Fig. S7.

After processing, results were further filtered for potential non-mRNAs by removing any genes with anomalously high read counts compared to other genes in the genome. RPKM values were calculated using gene lengths and total read counts from trimmed metatranscriptomic samples, as above. Each gene was then assigned a percentile expression value using the *stats* package in Python. Percentile gene expression / RPKM values were then visualized on a bulk and per-genome basis according to their functional annotation and/or taxonomic affiliation.

## Ethics approval and consent to participate

Not applicable

## Consent for publication

Not applicable

## Availability of data and materials

Newly-resolved MAGs used in this study as well as raw metagenomic reads from the OC1703A transect are available through NCBI at PRJNA1054206. Source information for all genomes is listed in Supplementary Table S2. Accession numbers for all public sequencing reads used for abundance and gene expression analyses are listed in Table S8/S10.

## Authors’ contributions

A.L.J. and A.E.D. designed the project. A.E.D. collected samples. A.L.J. and R.S.R.S. processed the metagenomic data and performed downstream bioinformatic analysis. A.L.J. and A.E.D. interpreted data and wrote the manuscript. All authors made comments on the manuscript.

## Acknowledgements

We thank Zhichao Zhou, Xin Sun, Chris Greening, Rachel Lappan, Alexa Nicolas, Noam Prywes, and Graciela Chavez for helpful discussions and data access. We also thank Alex Crits-Christoph for sharing code that was adapted for our metatranscriptomic analyses, and Alma Parada, Julian Fortney, and Nestor Arandia-Gorostidi for collecting and/or processing samples used for the metagenomic analysis. We thank the captain, crew, and science party of the R/V *Oceanus* OC1703A expedition. Finally, we acknowledge the Stanford Research Computing Center for informatic support as well as the Stanford Geomicrobiology Shared Laboratories Core Facility (RRID:SCR_025000).

## Supplementary Figures

**Figure S1.**
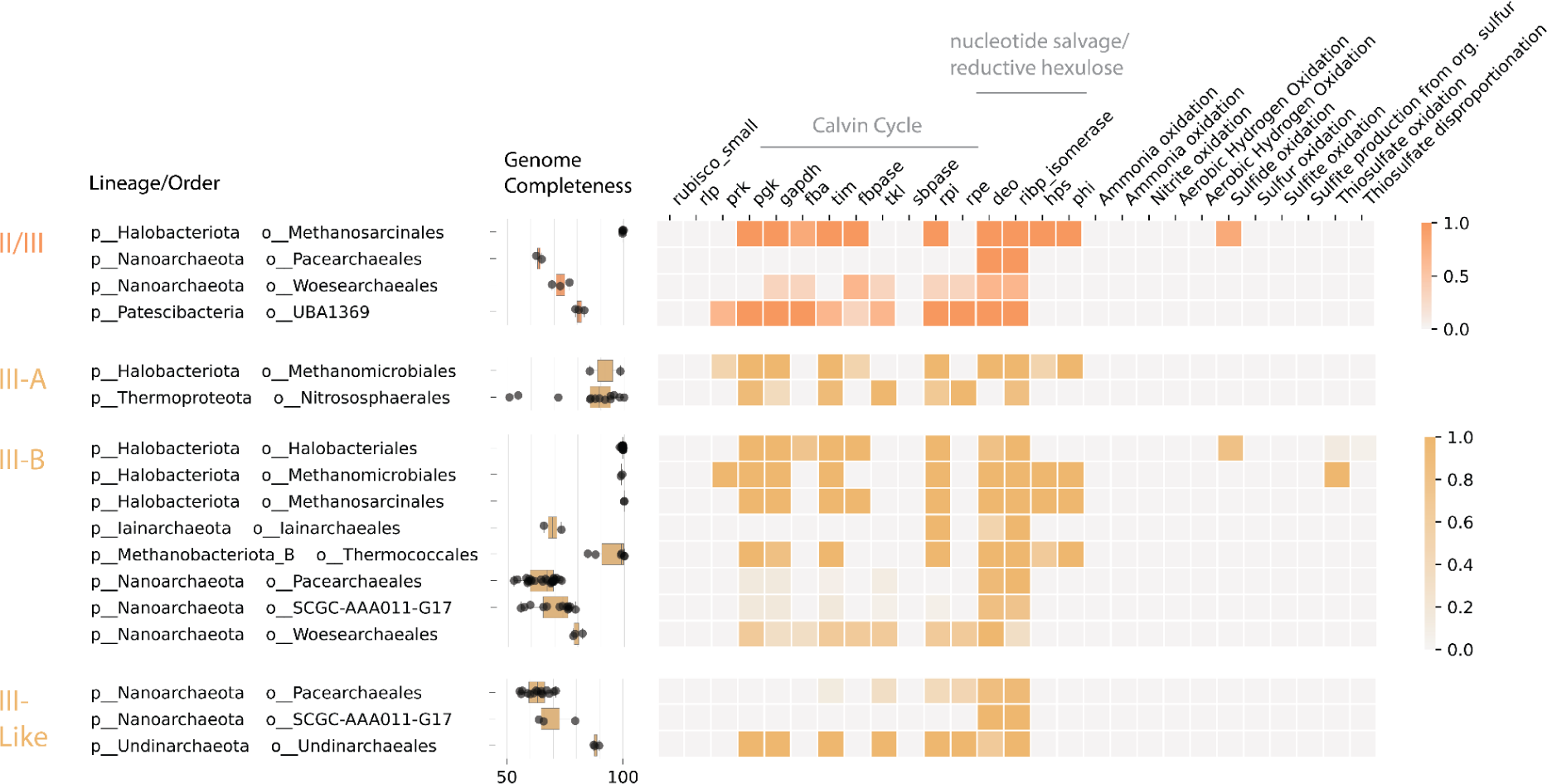
Characteristics of rubisco-encoding organisms (REOs) with the form II/III or form III-related enzyme types. Organisms were clustered into species groups (95% ANI) and were aggregated at the order level. **a)** For each order-level lineage, the completeness of representative genomes for each species group, and fraction of representative genomes that encode various rubisco-associated genes as well as those involved in oxidation of sulfur or nitrogen compounds for energy gain. Only those order-level lineages with more than one species group are displayed.

**Figure S2.**
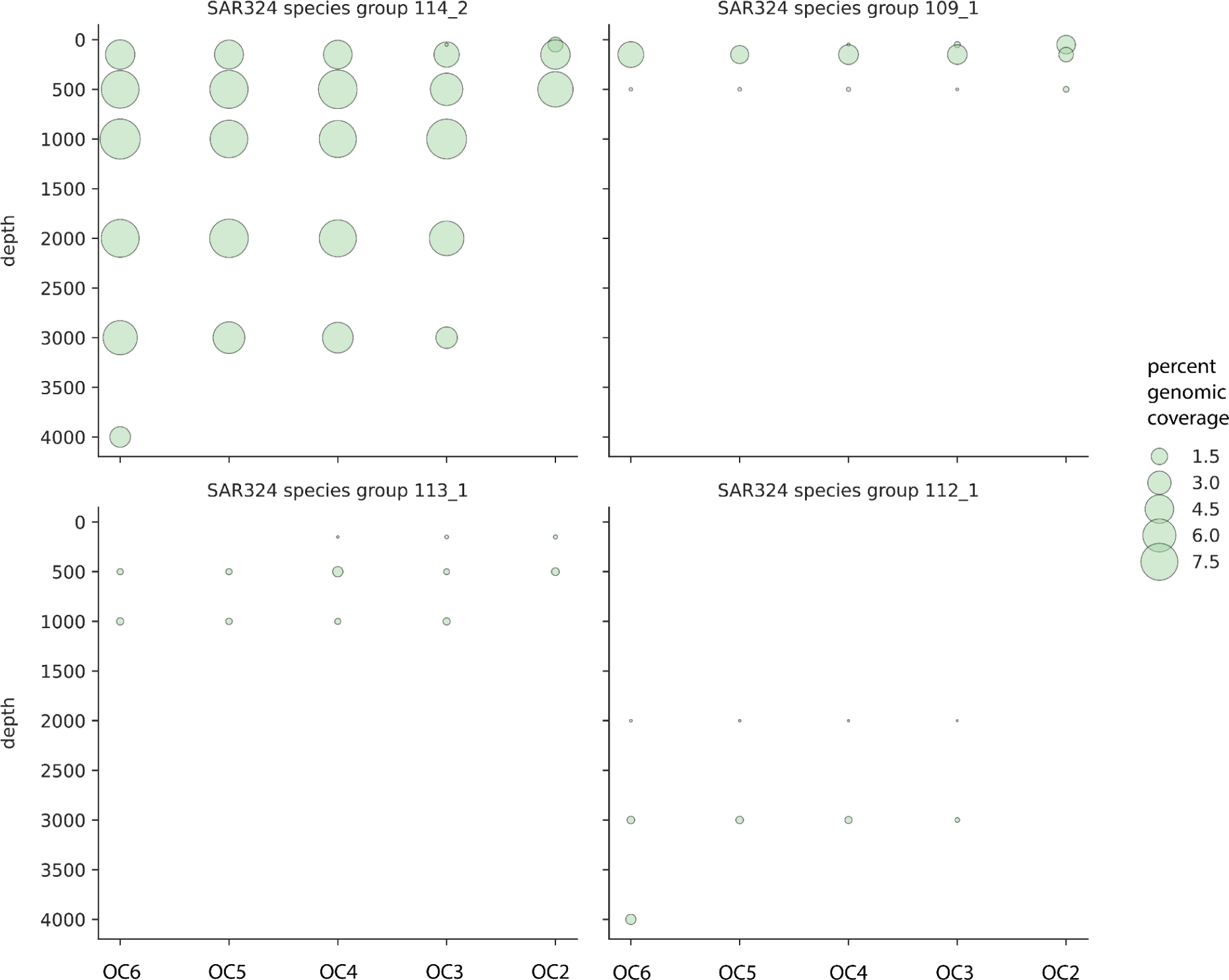
Distribution and abundance of SAR324 species groups with the form I rubisco across the OC1703A transect (coastal California). Dotplots represent 5 sites (OC2-6) over a span of 300 km from shore sampled across depths. Size of dots represent the relative abundance (calculated as the percentage of total genomic sequencing coverage).

**Figure S3.**
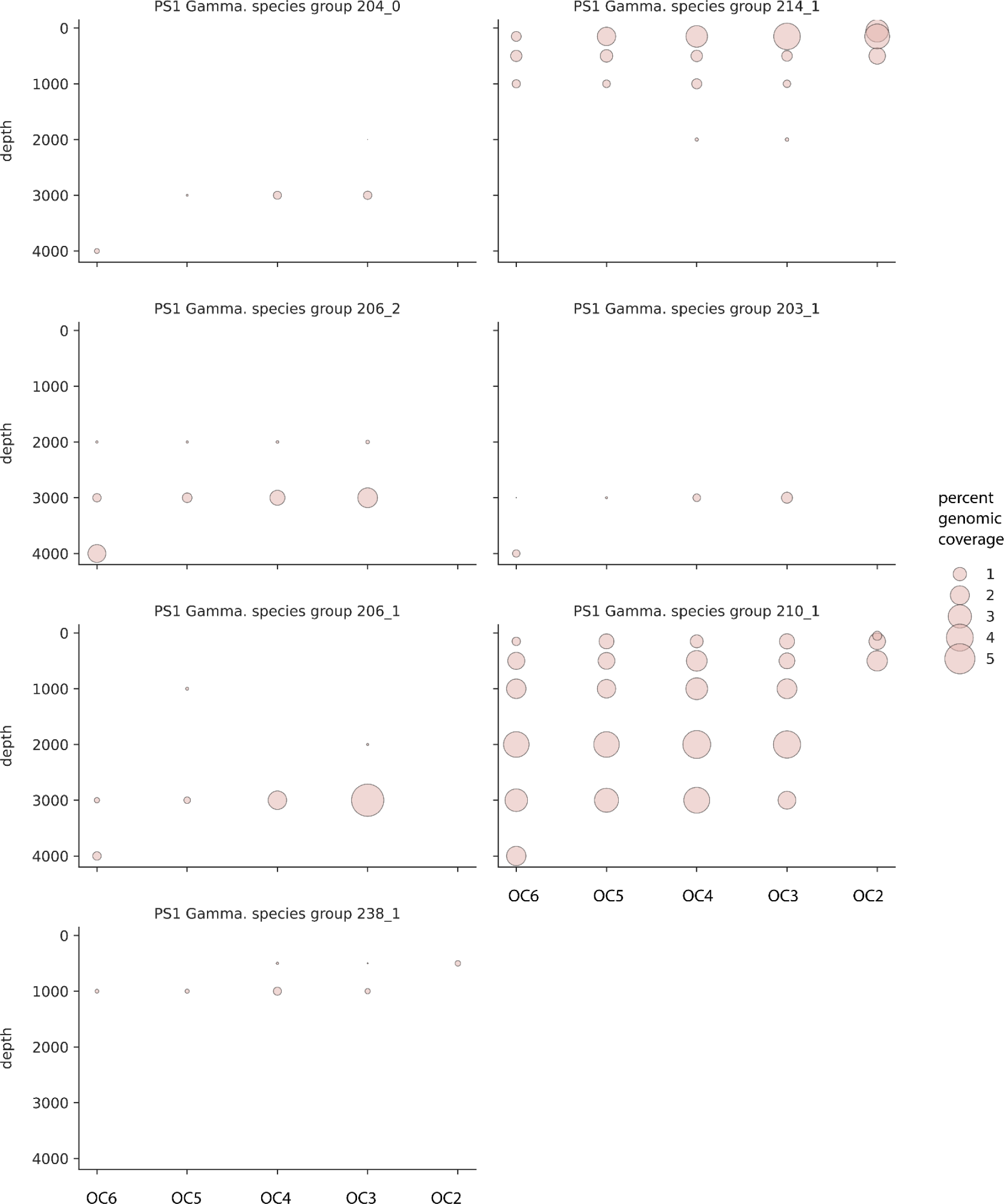
Distribution and abundance of PS1 (Gammaproteobacteria) species groups with the form II rubisco across the OC1703A transect (coastal California). Dotplots represent 5 sites (OC2-6) over a span of 300 km from shore sampled across depths. Size of dots represent the relative abundance (calculated as the percentage of total genomic sequencing coverage).

**Figure S4.**
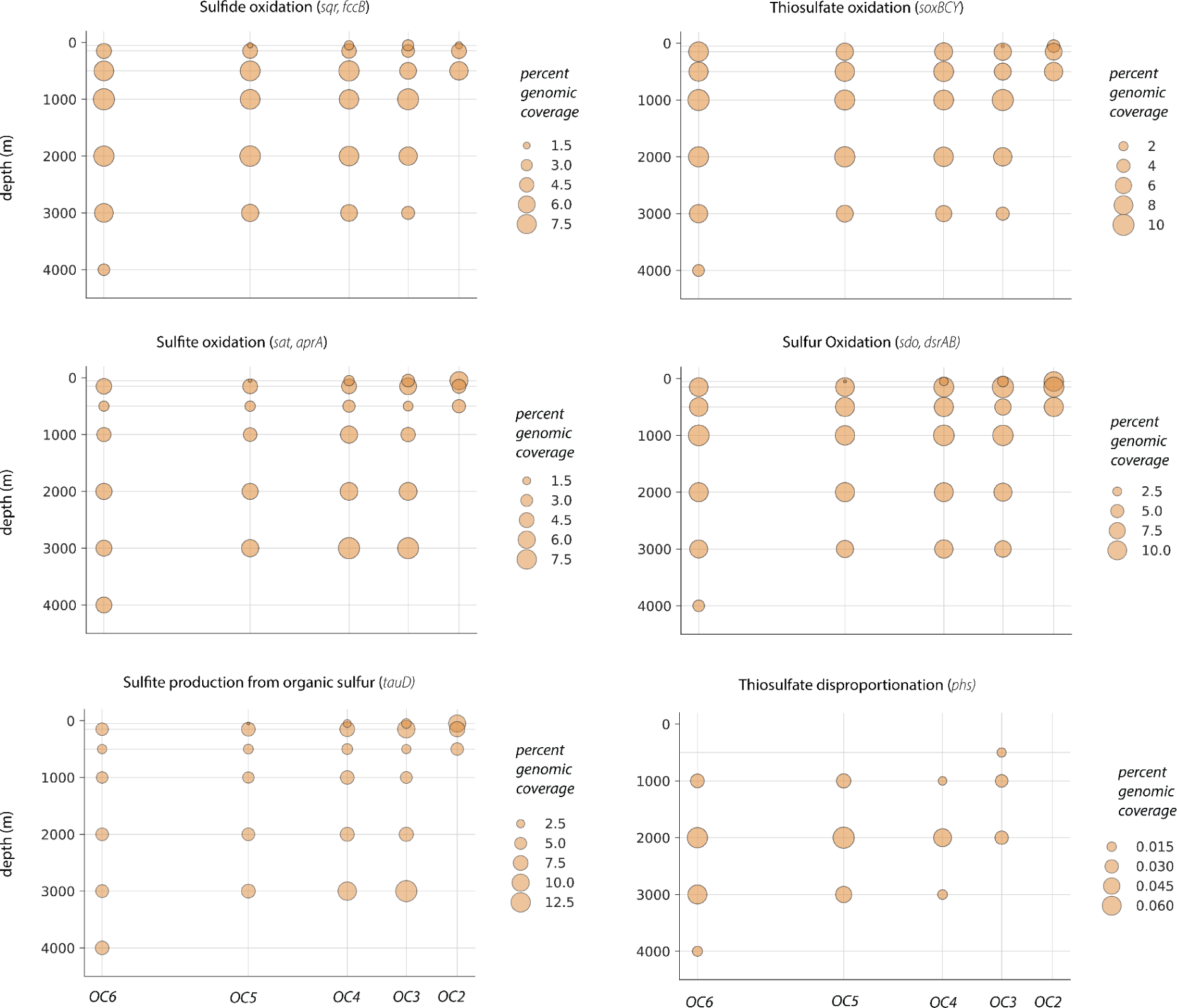
Distribution and abundance of REOs with the genetic capacity for various sulfur oxidation pathways across the OC1703A transect (coastal California). Dotplots represent 5 sites (OC2-6) over a span of 300 km from shore sampled across depths. Size of dots represent the summed relative abundance (calculated as the percentage of total genomic sequencing coverage) of all organisms encoding a given gene/pathway.

**Figure S5.**
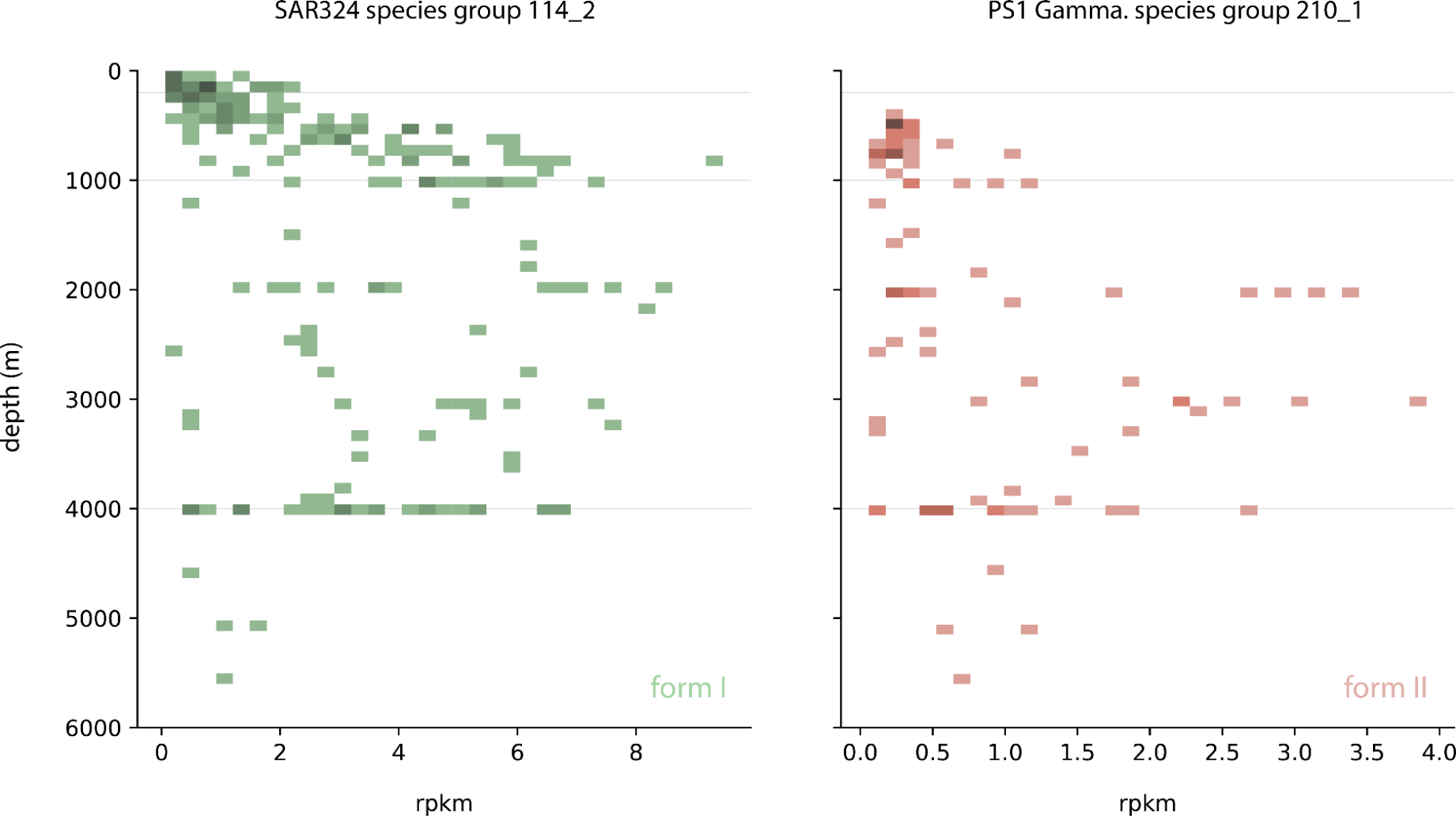
Global distribution and abundance of two rubisco-encoding species groups from the SAR324 and PS1 Gammaproteobacteria orders, respectively. Relative abundance is expressed as RPKM (reads per kilobase million). Shaded cells indicate the detection of one or more organisms in a given depth/RPKM bin, with hue intensity indicating the density of observations in that bin.

**Figure S6.**
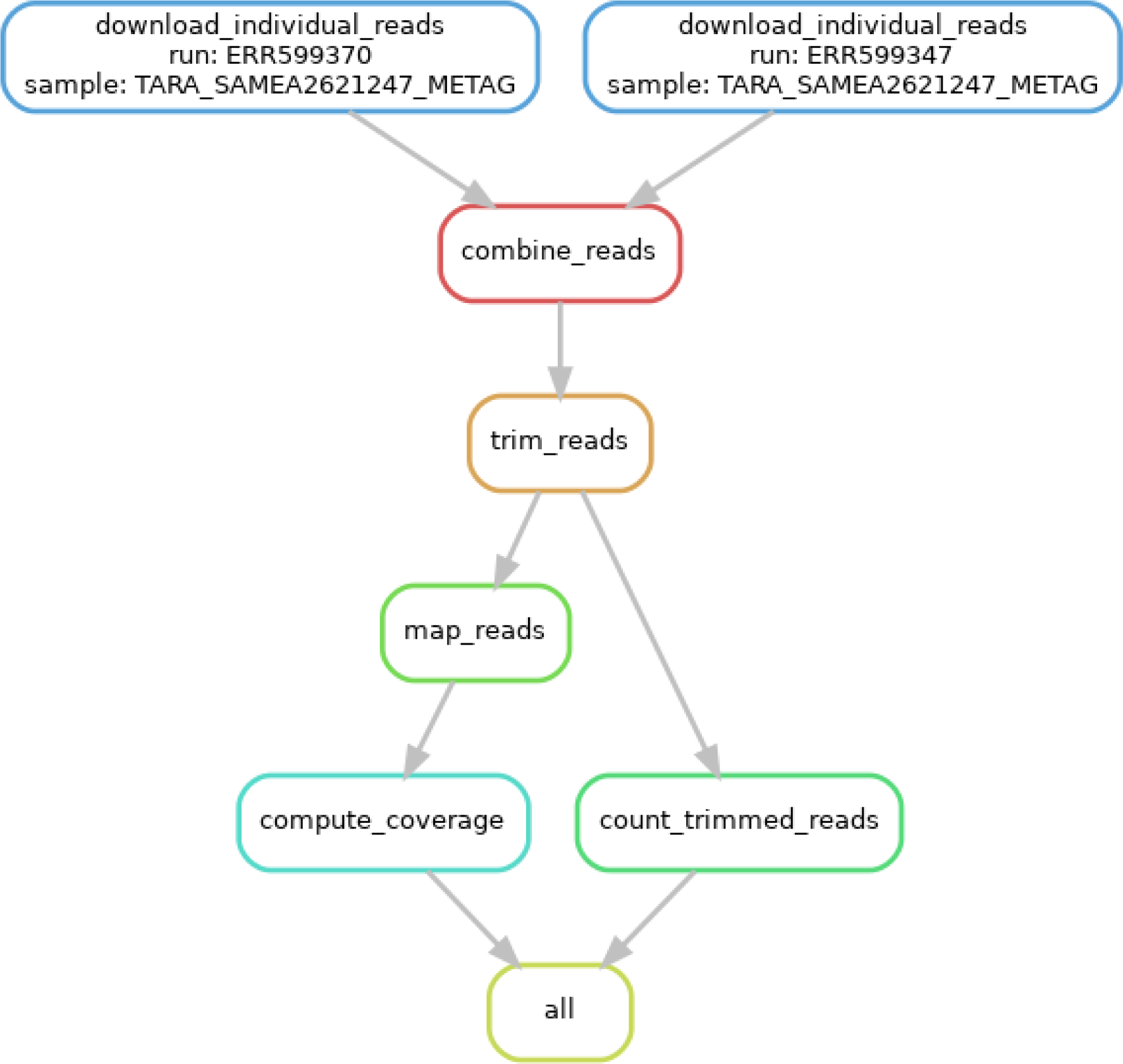
Conceptual overview of the computational workflow (implemented in Snakemake) used to align reads from global water column metagenomes to the REO genome set.

**Figure S7.**
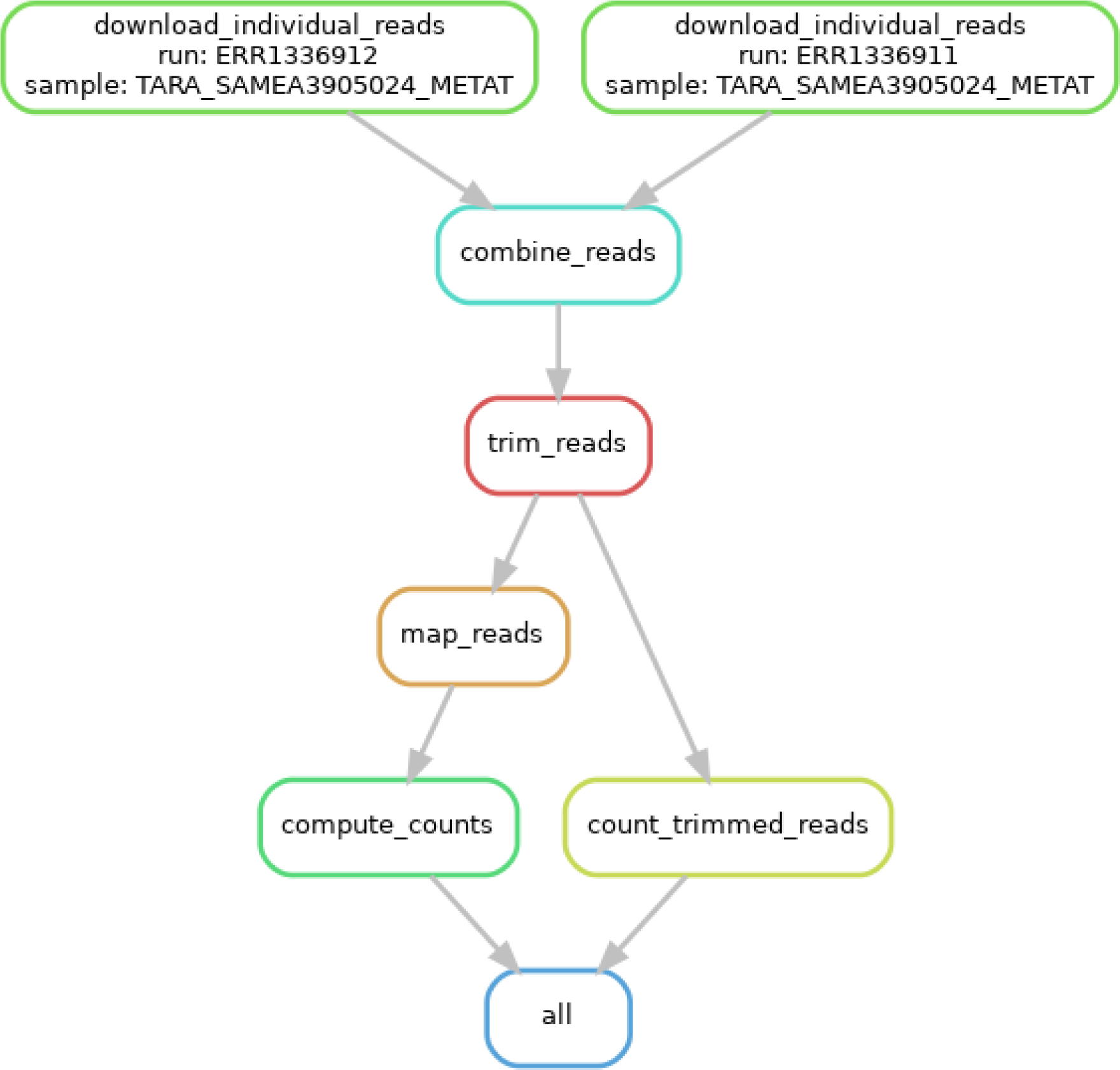
Conceptual overview of the computational workflow (implemented in Snakemake) used to align reads from global water column metatranscriptomes to the REO genome set.

